# Amygdala-cortical control of striatal plasticity drives the acquisition of goal-directed action

**DOI:** 10.1101/2020.02.28.970616

**Authors:** Simon D. Fisher, Lachlan A. Ferguson, Jesus Bertran-Gonzalez, Bernard W. Balleine

**Affiliations:** School of Biomedical Sciences and Pharmacy, University of Newcastle, Newcastle, NSW 2308, Australia; Decision Neuroscience Lab, School of Psychology, UNSW Sydney, NSW 2052, Australia

**Keywords:** Goal-directed learning, corticostriatal plasticity, intratelencephalic neurons, basolateral amygdala, prelimbic cortex, mouse

## Abstract

The acquisition of goal-directed action requires the encoding of specific action-outcome associations involving plasticity in the posterior dorsomedial striatum (pDMS). We first investigated the relative involvement of the major inputs to the pDMS argued to be involved in this learning-related plasticity, from prelimbic prefrontal cortex (PL) and from the basolateral amygdala (BLA). Using ex vivo optogenetic stimulation of PL or BLA terminals in pDMS, we found that goal-directed learning potentiated the PL input to direct pathway spiny projection neurons (dSPNs) bilaterally but not to indirect pathway neurons (iSPNs). In contrast, learning-related plasticity was not observed in the direct BLA-pDMS pathway. Using toxicogenetics, we ablated BLA projections to either pDMS or PL and found that only the latter was necessary for goal-directed learning. Importantly, transient inactivation of the BLA during goal-directed learning prevented the PL-pDMS potentiation of dSPNs, establishing that the BLA input to the PL is necessary for the corticostriatal plasticity underlying goal-directed learning.

## Introduction

The ability of humans and other animals to engage in goal-directed action has been found to rely on their capacity to encode the relationship between actions and their consequences or outcome, and to integrate that information with the current value of the outcome (Balleine and Dickinson, 1998; Dickinson and Balleine, 1994). Furthermore, recent evidence suggests that the acquisition of goal-directed actions relies on a cortical-limbic-striatal circuit centered on the posterior dorsomedial striatum (pDMS; Balleine, 2019; Balleine and O’Doherty, 2009). Inputs to the pDMS from prelimbic cortex (PL; Hart et al., 2018b, 2018a) and the basolateral amygdala (BLA; Balleine et al., 2003; Corbit et al., 2013) have been argued to be particularly important for this form of learning, although their relative importance in the cellular changes underlying the specific plasticity processes that control such learning is unknown.

Within the striatum, two classes of principal neuron have been described based on their projections to basal ganglia output nuclei and expression of dopamine receptors: direct pathway spiny projection neurons (dSPNs) primarily project directly to the substantia nigra pars reticulata (SNr) and express dopamine D1 receptors; and indirect pathway spiny projection neurons (iSPNs) primarily project indirectly to the SNr via the globus pallidus externa (GPe) and express dopamine D2 receptors. Importantly, evidence of learning-related plasticity has previously been documented at both of these classes of neuron (Kreitzer and Malenka, 2008; Lovinger, 2010) and, indeed, has been found to be induced specifically by the acquisition of goal-directed directed action (Shan et al., 2014). Nevertheless, in addition to the relative involvement of the PL and BLA pathways in pDMS plasticity, it remains to be resolved whether plasticity in these pathways is specific to inputs to dSPNs, iSPNs or to both of these cell types.

In this study, we used *ex vivo* optogenetic stimulation of PL and BLA terminals in the pDMS to assess learning-related plasticity selectively at these inputs in mice. Using behavioral methods and *ex vivo* whole-cell recordings of dSPNs and iSPNs in the pDMS, we demonstrate goal-directed learning-related plasticity in both the ipsilateral and contralateral PL-pDMS pathway and so, confirming recent reports (Hart et al., 2018a), that intratelencephalic neurons in the PL are critical for this effect. Furthermore, we found that these changes in plasticity were localized to dSPNs and not iSPNs. Importantly, changes in plasticity were not observed in the BLA-pDMS pathway. Nevertheless, we found that the direct BLA-PL projection was necessary for plasticity in the PL-pDMS pathway and, therefore, evidence that the BLA and PL function together to support goal-directed learning.

## Results

### Goal-directed learning potentiates prelimbic inputs to dSPNs in the pDMS

Prior attempts to investigate the role of inputs to the pDMS in plasticity induced by goal-directed learning have evoked potentials in pDMS by stimulating fibers in the corpus collosum. To directly assess plasticity in the PL-pDMS pathway, a pathway previously found to be essential for goal-directed learning (Hart et al., 2018b), we infused channelrhodopsin (ChR2) into the PL and used optogenetic stimulation at PL terminals in the pDMS to evoke potentials in the PL-pDMS pathway *ex vivo*. We used Drd2-EGFP mice injected unilaterally with ChR2-mCherry in the PL (**Fig. 1A-C; Fig. S1**) that, following recovery, were divided into two groups; one trained to lever press for pellets (**Fig. 1D**), and a second yoked control group that received the lever press and reward delivery unpaired. 24 hours after the final training session mice were processed for *ex vivo* electrophysiological recordings of striatal slices. Whole field 470 nm light stimulation of PL terminals elicited postsynaptic potentials and action potentials in D2-GFP positive iSPNs, and D2-GFP negative putative dSPNs in the pDMS under current-clamp (**Fig. 1E-F**). Under voltage-clamp recordings, excitatory postsynaptic currents (EPSCs) were elicited by brief optical stimulation at −70 mV and +40 mV to measure AMPA and NMDA components respectively (**Fig. 1G**). An increase in AMPA current has been found to reliably mediate long-term potentiation (Malinow and Malenka, 2002). The ratio between AMPA and NMDA currents was used as the measure of postsynaptic synaptic plasticity, as this measure is independent of the number of synapses activated and therefore independent of inherent variables such as ChR2 virus expression and light dispersion through the tissue.

**Figure 1.**
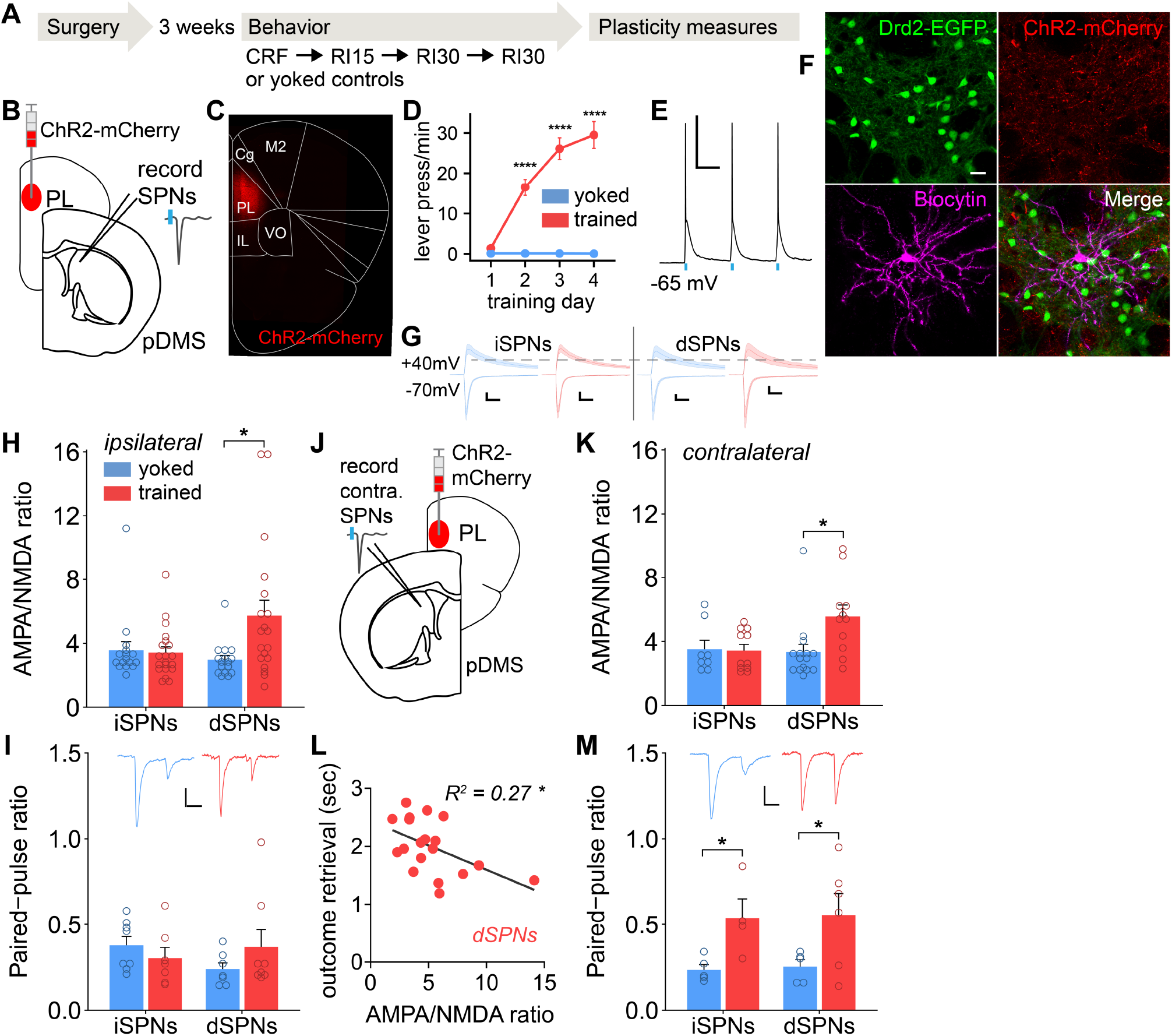
Goal-directed learning induces potentiation of the PL-pDMS pathway to dSPNs. **(A)** Experimental protocol across time. CRF = Continuous Reinforcement. RIX = Random Interval of X seconds schedule. (**B-C**) ChR2 was injected unilaterally into the PL of Drd2-EGFP mice. (**D**) Mice in the trained group (n=12), with reward delivery contingent on lever presses, had a significantly higher press rate than yoked controls (n=9) for which reward delivery was non-contingent (two-way RM ANOVA: session x group, F_(3,57)_ = 33.59, *P* < 0.0001; post hoc tests, \*\*\*\**P* < 0.0001). (**E**) Under current-clamp, 10 ms pulses of 470 nm light elicited action potentials in recorded SPNs. Scale bars = 100 ms and 40 mV. (**F**) Whole cell, patch-clamp recordings were made from SPNs in the pDMS of Drd2-EGFP mice. Fibers from BLA neurons infected with ChR2-mCherry were visible. The pipette was filled with biocytin to label recorded cells. Scale bar = 20 um respectively. (**G-H**) In the ipsilateral PL-pDMS pathway, the AMPA/NMDA ratio did not change due to learning (trained = red, control = blue) in iSPNs (two-way ANOVA: behavior x cell type, F_(1,68)_ = 5.5, *P* = 0.02; post hoc test, *P* = 0.99), and was significantly greater in dSPNs (*P* = 0.01). Scale bar = 25pA, 25ms. (**I**) paired-pulse ratio measures were not affected by training in iSPNs or dSPNs (two-way ANOVA: behavior, F_(1,26)_ = 0.58, *P* = 0.45). Inset shows representative paired-pulse traces. **(J)** Whole cell, patch-clamp recordings were made from SPNs in the contralateral pDMS (to the PL injection site) of Drd2-EGFP mice. (**K**) The AMPA/NMDA ratio did not change due to learning in iSPNs (two-way ANOVA: behavior x cell type, F_(1,41)_ = 4.17, *P* = 0.04; post hoc test, *P* = 1.0), and was significantly greater in dSPNs (*P* = 0.02). (**L**) A significant correlation was found between AMPA/NMDA ratios from dSPN recordings and the mean outcome retrieval time of the first 10 outcomes during the RI15 training session of the corresponding mouse (Pearson R^2^ = 0.27, two-tailed *P* = 0.024). (**M**) Facilitation of paired-pulse ratio measures was found in projections on to both iSPNs and dSPNs as a result of training (two-way ANOVA: behavior, F_(1,16)_ = 4.92, *P* = 0.04).

We found that PL projections to iSPNs exhibited no difference in the AMPA/NMDA ratio in trained animals that learned the action-outcome association relative to the yoked controls that did not (**Fig. 1H**). In contrast, PL projections on to dSPNs were significantly potentiated in trained versus control animals following the acquisition of goal-directed action. No differences in the paired-pulse ratio, a measure of presynaptic plasticity, were found in PL projections on to either SPN cell population (**Fig. 1I**). To rule out the possibility that AMPA/NMDA ratio differences were due to higher levels of task engagement or motor output in trained mice, entry rate into the food magazine and the time taken to collect food reward were compared between trained and yoked animals. The measures suggested that yoked mice were engaged and active comparably to trained mice – with significantly higher magazine entries per minute than trained animals, and only moderately longer time to collect rewards (**Figs. S2A and S2B**). These differences likely reflect the more uncertain reward environment that control mice were exposed to. Together, these results demonstrate that goal-directed learning induces potentiation in the PL-pDMS pathway, exclusively on to dSPNs.

### Intratelencephalic PL neurons support goal-directed PL-pDMS plasticity

Recent findings suggest that the bilaterally-projecting intratelencephalic (IT) PL neurons, as opposed to ipsilaterally-projecting pyramidal tract (PT) PL neurons, are critical for encoding the action-outcome associations necessary for goal-directed learning (Hart et al., 2018a). In measuring the ipsilateral PL-pDMS pathway, we likely stimulated a mixed population of IT and PT afferents. To determine if these plasticity findings also are present when IT projections from the PL were exclusively measured, the same recordings were performed in the pDMS contralateral to the PL ChR2 injection site (**Fig. 1J**). Matching the ipsilateral findings, contralateral PL projections on to iSPNs exhibited no difference in the AMPA/NMDA ratio, and projections on to dSPNs were significantly potentiated, following goal-directed learning (**Fig. 1K**). A similar proportion of dSPNs responded to optical stimulation in the ipsilateral and contralateral pathways (**Figs. S3A and S3B**). To investigate the relationship within subjects between plasticity measures and early learning performance, we correlated AMPA/NMDA ratios with the mean time taken to retrieve an earned outcome in the first 10 outcome deliveries of the first random interval session (RI15). A significant negative correlation was found between this outcome retrieval time and the AMPA/NMDA ratios from ipsi and contralateral dSPN recordings (**Fig. 1L**). Paired-pulse measures of the contralateral PL-pDMS pathway showed enhanced facilitation of projections on to both iSPNs and dSPNs due to goal-directed learning (**Fig. 1M**). Paired-pulse facilitation indicates decreased presynaptic release probability (Debanne et al., 1996; Kreitzer and Malenka, 2005), and hence likely reflects a different plasticity process than that of the postsynaptic potentiation found for projections on to dSPNs. These findings support prior findings that IT PL projections to the pDMS are critical for goal-directed learning.

### BLA projections to the pDMS are not modified by goal-directed learning

Multiple lines of evidence demonstrate that the BLA plays a critical role in goal-directed learning. Cell-body lesions of the BLA was reported to abolish sensitivity to outcome devaluation and contingency degradation after goal-directed training (Balleine et al., 2003; Ostlund and Balleine, 2008), as was disconnection of the BLA and pDMS using asymmetrical lesions of these structures (Corbit and Balleine, 2005; Corbit et al., 2013). Moreover, many studies have indicated that BLA afferents provide critical information related to the motivational significance of outcomes, which is required for goal-directed learning (Balleine and Killcross, 2006; Balleine et al., 2003; Corbit and Balleine, 2005; Corbit et al., 2013). To further probe BLA function, therefore, we measured learning-related plasticity in the BLA-pDMS pathway, using the experimental procedures previously described. Drd2-EGFP mice were unilaterally injected in the BLA with ChR2 and underwent goal-directed training or yoked control sessions (**Figs. 2A-C and S4**). We found that BLA projections on to dSPNs or iSPNs exhibited no difference in the AMPA/NMDA ratio following goal-directed learning (**Fig. 2D-F**). Additionally, no difference in the paired-pulse ratio was found due to training (**Fig. 2G**). Of healthy dSPNs recorded, 62% responded to BLA optical stimulation, which is comparable to the 66% of dSPNs that responded to ipsilateral PL optical stimulation (**Fig. S3A,C**). Of healthy iSPNs recorded, 93% responded to BLA optical stimulation (**Fig. S3C**). In sum, these findings indicate that the BLA-pDMS pathway does not undergo long-term plasticity as a result of goal-directed learning.

**Figure 2.**
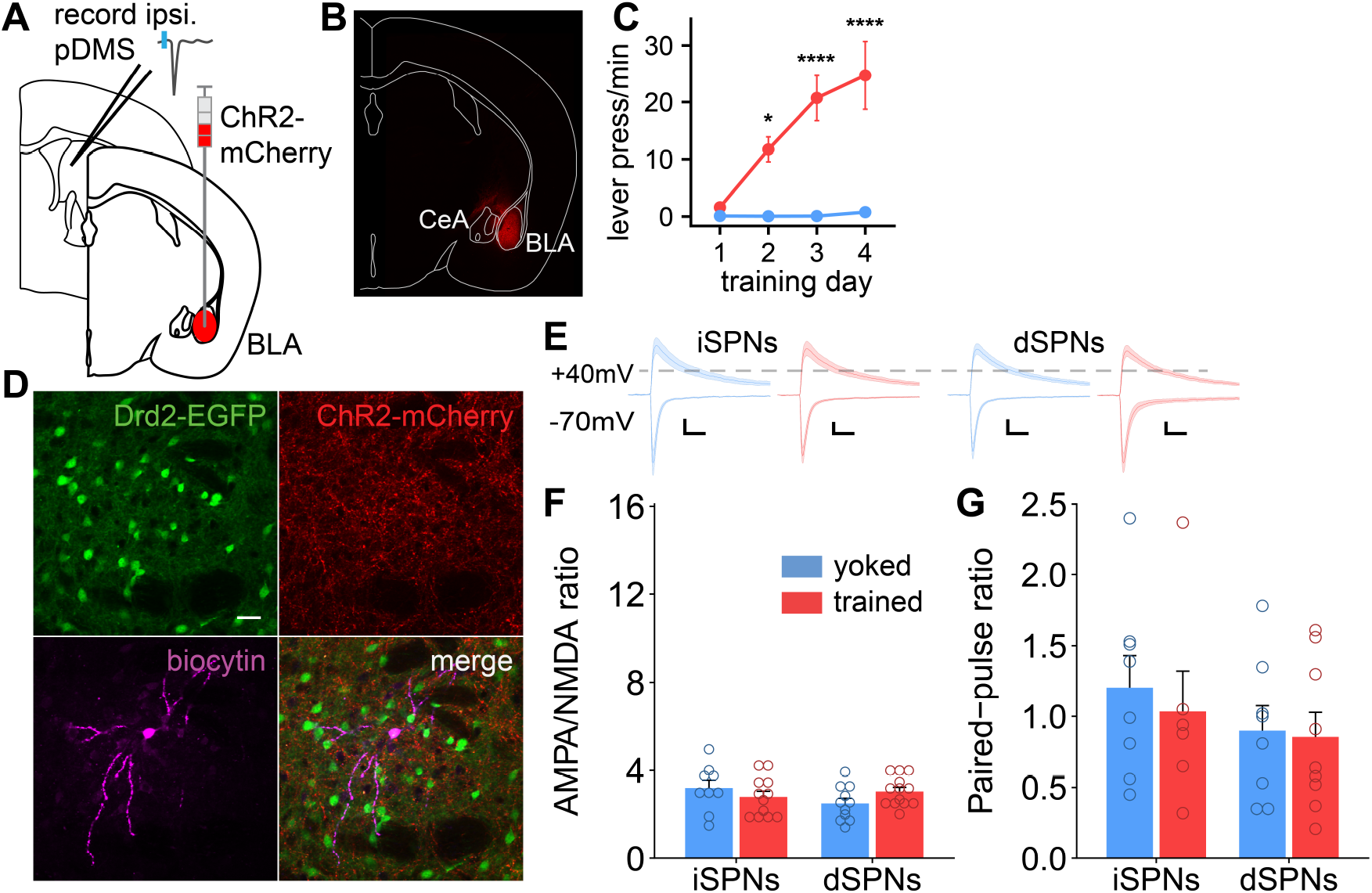
BLA projection to pDMS does not undergo plasticity following goal-directed learning. (**A-B**) ChR2 was injected unilaterally into the basolateral amygdala (BLA) of Drd2-EGFP mice. (**C**) Mice in the trained group (n=7), with reward delivery contingent on lever presses, had a significantly higher press rate than yoked controls (n=6) for which reward delivery was non-contingent (two-way RM ANOVA: session x group, F_(3,33)_ = 10.69, *P* < 0.0001; post hoc tests, \**P* < 0.05, \*\*\*\**P* < 0.0001). (**D**) Whole cell, patch-clamp recordings were made from SPNs in the pDMS of Drd2-EGFP mice. Fibers from BLA neurons infected with ChR2-mCherry were visible. The pipette was filled with biocytin to label recorded cells. Scale bar = 20 um. (**E-F**) In the ipsilateral BLA-pDMS pathway, the AMPA/NMDA ratio was not affected by learning in either SPN population (two-way ANOVA: behavior, F_(1,34)_ = 0.74, *P* = 0.4). Scale bar = 25pA, 25ms. (**G**) Paired-pulse ratio measures were not affected by training in iSPNs or dSPNs (two-way ANOVA: training, F_(1,27)_ = 0.29, *P* = 0.6).

### The direct BLA projection to the PL but not to pDMS is critical for goal-directed learning

If the direct BLA to pDMS projection is unaffected by goal-directed learning, the question arises what is the BLA contribution to this learning established by the previous studies described above. One possibility is that, rather than altering plasticity in the pDMS through the direct pDMS projection, the BLA contributes to this plasticity indirectly via its well documented projection to the PL (Gabbott et al., 2006; Groenewegen et al., 1997). To assess this question, we first used pathway-specific toxicogenetic ablation of BLA projection neurons coupled with a behavioral assessment of goal-directed action control using outcome devaluation procedures (**Fig. 3A-D**). Adult wild-type mice were injected with a retro-Cre virus into either the PL or the pDMS, and a Cre-dependent Caspase3 virus into the BLA, thereby lesioning BLA neurons projecting directly to either the PL or pDMS, respectively (**Fig. S5**). Control animals received the retro-Cre virus infused into either the PL or pDMS, plus an infusion of PBS into the BLA. We subsequently found no differences between these infusion conditions (largest F_(1,6)_ = 3.617, *P* = 0.11) and so combined them into a single control group.

**Figure 3.**
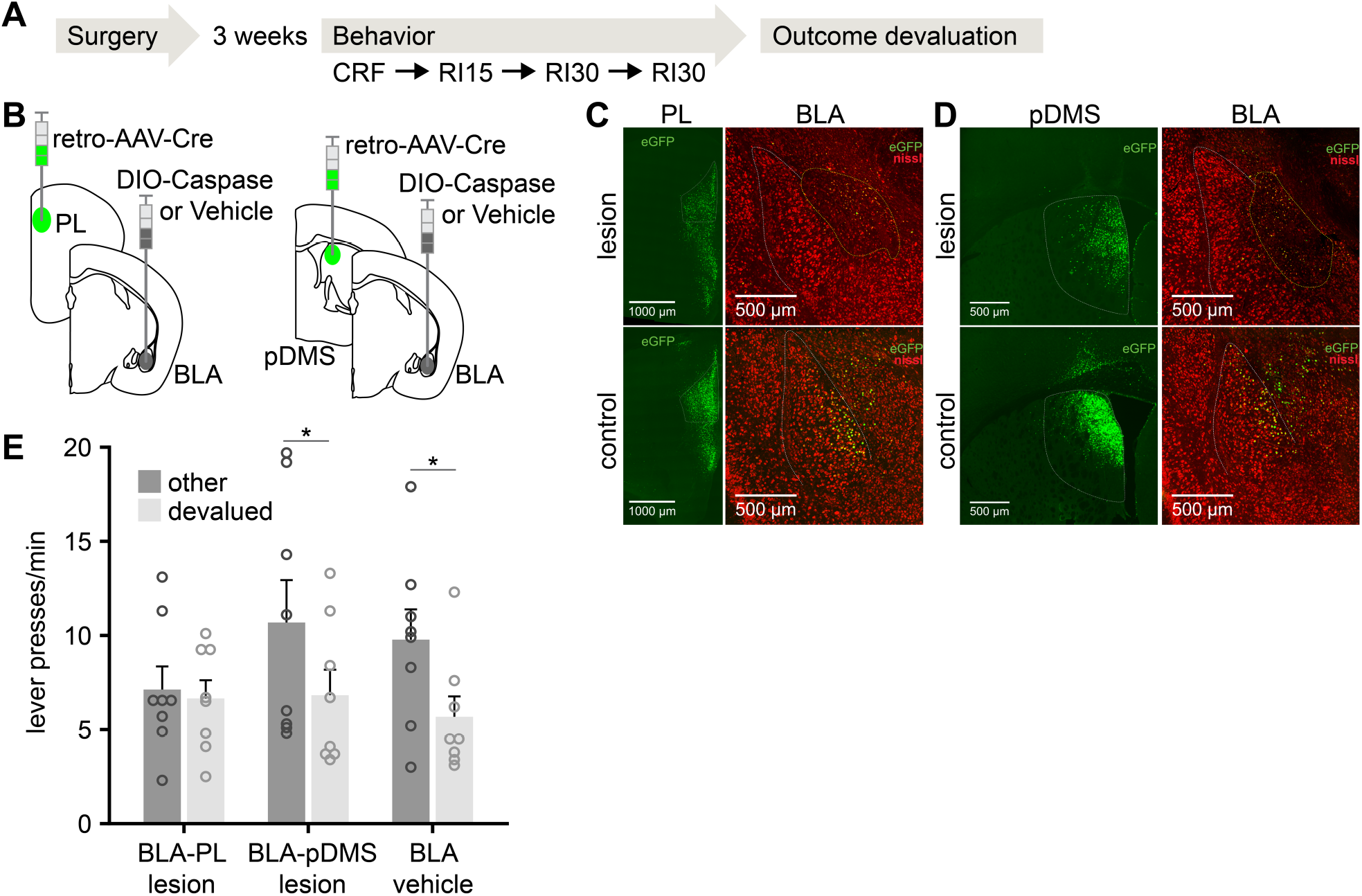
BLA projection to prelimbic cortex, and not pDMS, supports goal-directed learning. (**A**) Experimental protocol, where CRF = Continuous Reinforcement, and RIX = Random Interval of X seconds schedule. (**B**) C57BL/6 mice were injected bilaterally with retro-AAV-Cre-GFP in either the PL (n=8) or the pDMS (n=8), and Cre-dependent DIO-Caspase in the BLA. The vehicle control group (n=12) were injected with retro-AAV-Cre in either the PL or pDMS, and PBS in the BLA. Retrograde transport of the retro-AAV-Cre-GFP construct from the PL (**C**) and the pDMS (**D**) to the BLA was observed, and in the ‘lesion’ group the Cre-dependent Caspase construct induced cell death as seen with reduced visualization of nissl bodies. (**E**) Mice with a specific lesion of the BLA-PL pathway were insensitive to outcome devaluation, hence not responding significantly less for the devalued outcome (F_(1,19)_ = 0.01, *P* > 0.05). In contrast, mice with BLA-pDMS pathway lesions (F_(1,19)_ = 4.97, *P* < 0.05) and control animals (F_(1,19)_ = 5.60, *P* < 0.05) remained sensitive to outcome devaluation.

After recovery, mice were food-deprived and trained to press a lever for a food pellet reward (**Fig. 3A**) after which the degree to which their lever press performance was goal-directed was assessed using two outcome devaluation tests (Shan et al., 2014); one conducted after satiety on the pellets and one after satiety on the maintenance diet (counterbalanced). These tests revealed that, whereas mice in the control group and the group given a lesion of the BLA neurons projecting directly to the pDMS showed a reliable devaluation effect – pressing less on test after satiety on the pellet outcome relative to satiety on their maintenance diet – mice in the group in which BLA neurons projecting to the PL were lesioned failed to show this effect, pressing similarly in both tests (**Fig. 3E**). These findings demonstrate that with both pathways similarly lesioned, whereas the BLA’s direct input to the pDMS is not necessary for goal-directed learning, its input to the PL is essential; mice with lesions of the BLA-PL pathway were impaired in forming the action-outcome associations necessary to acquire goal-directed actions, suggesting the BLA involvement in this acquisition process is achieved via its projection to the PL area.

### Inhibition of BLA neurons during learning abolishes PL-pDMS plasticity

The current findings, together with prior work, suggest that, while the BLA is involved in goal-directed learning, the BLA-pDMS projection is not critical for this learning. It could be the case that the BLA functions via projections to the PL to support plasticity in the PL-pDMS pathway found critical for goal-directed learning. To test this hypothesis mice were injected with ChR2 in the PL as previously described, and additionally with either the inhibitory hM4D designer receptor exclusively activated by designer drugs (DREADD) or mCherry control virus in the BLA (**Fig. 4A-D**). The efficacy of the hM4D virus was confirmed *in vitro* with current-clamp recordings of BLA neurons, demonstrating the inhibitory effect (**Fig. 4E-F**) of CNO application on these neurons. Both the hM4D and the mCherry control groups were trained on the contingent version of the goal-directed learning task. Additionally, all mice from both groups were injected with CNO 40 minutes prior to the session with the aim of inhibiting the BLA in the hM4D, but not mCherry, mice. During goal-directed learning, there was no difference in task performance between mice injected with hM4D and mCherry controls, until the final training session, in which hM4D mice exhibited an unexpected reduction in pressing rate (**Fig. 4G**). However, these animals did not differ from the control group in other performance measures, including magazine entry rates and the time to collect reward (**Fig. S6**). Inhibition of BLA neurons during training (hM4d group) significantly reduced the AMPA/NMDA ratio in the PL-pDMS pathway on to dSPNs, in comparison with mCherry controls (**Fig. 4H-I**). The AMPA/NMDA ratio of the hM4D group was not significantly different to the ratio previously found with yoked controls that did not learn the action-outcome relationship (one-way ANOVA, F_1,28_ = 0.52, P = 0.48). Additionally, the AMPA/NMDA ratio of the control mCherry group was not significantly different to that found previously in dSPNs with trained animals (one-way ANOVA, F_1,38_ = 2.33, P = 0.14). The PL-pDMS paired-pulse ratio was not altered by inhibition of BLA neurons (**Fig. 4J**), consistent with the lack of change seen in the prior PL-pDMS results between trained and yoked mice (Fig. 1J). Plasticity of iSPNs was not measured as this did not change with learning (**Fig. 2I**). These data demonstrate that inhibition of the BLA during learning abolished goal-directed learning-related plasticity in the PL-pDMS pathway.

**Figure 4.**
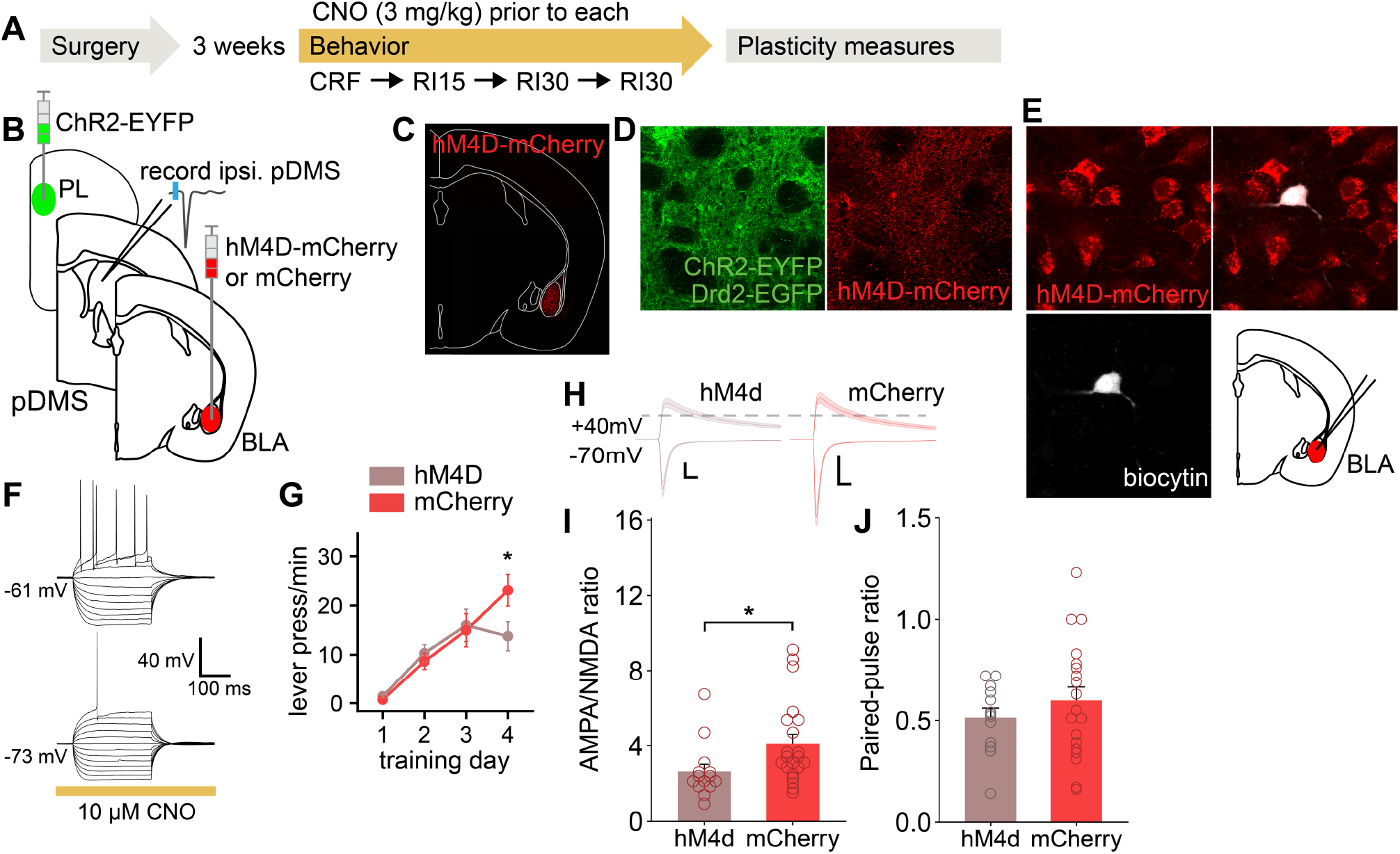
Plasticity of the PL projection to dSPNs in pDMS is impaired by BLA inactivation. (**A**) Experimental protocol was identical to previous groups (Fig. 1A) except all mice were injected with CNO (3 mg/kg i.p.) 40 minutes prior to each training session. (**B-C**) ChR2 was injected unilaterally into the PL, and either the inhibitory hM4D DREADD (n=6) or a mCherry control virus (n=6) was injected into the ipsilateral BLA. (**D**) In the pDMS, projections from infected PL cells and the endogenous Drd2-EGFP processes intermixed. Fibers from infected BLA neurons were also present. Scale bars = 1 mm and 20 um respectively. (**E-F**) Current-clamp recordings of infected BLA neurons demonstrated that CNO (10 uM) reliably hyperpolarized the membrane potential and increased the threshold required for action potentials. (**G**) Mice injected with hM4D and mCherry both learned the lever pressing task similar to prior groups. There was a significant interaction between virus and experiment session, with hM4D-injected mice pressing significantly less (two-way RM ANOVA: session x virus, F_(3,27)_ = 3.8, *P* = 0.02; post hoc tests, \**P* = 0.04). (**H-I**) The AMPA/NMDA ratio of the ipsilateral PL projections to dSPNs following learning was significantly reduced in mice injected in the BLA with the active hM4D DREADD in comparison to mCherry controls (oneway ANOVA, F_(1,33)_ = 4.78, \**P* = 0.03). Scale bar = 200pA, 25ms. (**J**) paired-pulse ratio measures did not differ by virus (one-way ANOVA, F_(1,30)_ = 0.89, *P* = 0.35).

## Discussion

Our results demonstrate that the PL projection to the pDMS plays a critical role in goal-directed learning, leading to significant potentiation of both the ipsilateral and contralateral input to dSPNs, but not iSPNs, in the pDMS. This plasticity is most likely postsynaptic, enabled by changes in AMPA or NMDA-mediated currents. In contrast, despite prior reports that the direct BLA inputs to the pDMS may play a role in goal-directed learning, we found that the BLA-pDMS projection does not undergo long-term plasticity as a result of this learning. We reasoned that one means by which the BLA could remain critical to goal-directed learning and associated plasticity in the pDMS is via its well described input to the PL. And, indeed, we found that toxicogenetic ablation of the PL-BLA pathway abolished the capacity for acquiring goal-directed learning in mice and, furthermore, that when the BLA was inhibited during that learning the potentiation in the PL-pDMS pathway was abolished.

### PL-pDMS pathway plays a critical role in goal-directed learning

That the changes in the PL-pDMS pathway persisted for at least 24 hours after training suggests that they reflect long-term potentiation (LTP) as a result of the goal-directed learning. This potentiation of PL-pDMS projections may account for the broad potentiation seen following the same behavioral task in prior work from our group (Shan et al., 2014), in which non-specific afferents on to SPNs in the pDMS (likely involving many cortical and potentially thalamic afferents) were electrically stimulated. Due to difficulties in comparing magnitudes of AMPA-NMDA ratio changes between studies that use broad electrical and pathway-specific optical stimulation of afferents (as used here), it is not possible to determine if PL-pDMS plasticity accounts for all of the general potentiation found previously, and hence it is possible that other pathways also undergo plasticity. The selective importance of the PL projection to goal-directed learning is, however, strongly supported by prior findings (e.g., Hart et al., 2018a; reviewed in Balleine, 2019), some of which have also characterized an important temporal limitation to PL involvement; i.e., pre-but not post-training manipulations of PL affect goal-directed learning, suggesting it plays only a transient role, with information required for expression of this underlying knowledge stored elsewhere (Ostlund and Balleine, 2005; Tran-Tu-Yen et al., 2009).

That area appears now to be the pDMS, due to its requirement to be intact for the expression of goal-directed learning (Yin et al., 2005), and the strong projections to the DMS from the PL (Berendse et al., 1992; McGeorge and Faull, 1989). Furthermore, feedback circuits from the striatum to the cortex, likely via thalamocortical projections (Cruikshank et al., 2012; Little and Carter, 2012), provide a pathway for striatal learning to influence future learning involving the PL. Indeed, lesions of the mediodorsal thalamus, a likely hub for this feedback (Parent and Hazrati, 1995), disrupt outcome devaluation performance and contingency degradation learning (Corbit et al., 2003). Thus, the PL-pDMS plasticity found in the current study would underlie critical, likely early, aspects of goal-directed learning that feed into a broader pDMS-based network.

### Plasticity on to direct pathway SPNs is critical for goal-directed learning

Given the critical involvement of the PL-pDMS pathway, one of the more important observations in the current series is, therefore, the finding of potentiation in the PL input to pDMS exclusively on to dSPNs, advancing prior work (Shan et al., 2014). The potentiation results presented here are also consistent with intracellular changes in dSPNs associated with goal-directed learning, including enhanced pMAPK expression in pDMS SPNs (Shan et al., 2014), which has demonstrated to be required for learning (Shiflett et al., 2010) and of other transcriptional events secondary to pMAPK such as histone phosphorylation (Matamales et al., 2020). Inhibiting the PL during learning prevents this increased phosphorylation (Hart et al., 2018b), further supporting the criticality of the PL input in the dSPN learning-related plasticity process. This change in the PL-pDMS pathway was specific to dSPNs; although prior work found depression of the global synaptic response on to iSPNs, no change on to these cells was found in the present study from either PL or BLA inputs.

### Intratelencephalic PL neurons underlie goal-directed learning

We also observed that learning-related plasticity in the contralateral PL-pDMS projection matched plasticity found in ipsilateral projection consistent with recent work demonstrating that goal-directed learning relies on bilaterally-projecting PL neurons. These crossed projections are generated by intratelencephalic (IT) neurons (Hart et al., 2018a) and suggest that the specific action-outcome associations underlying goal-directed learning are propagated bilaterally within the striatum. Although one might anticipate such propagation to be lateralized depending on the effector used for the specific action, in fact bilateral propagation of this information makes sense. The goal of goal-directed actions is to achieve a specific change in the world as opposed to a specific motor movement; for example, a hungry mouse is attempting to enhance food delivery and knowing that pressing the left lever gives it, say, food pellets will, therefore, promote the pressing of that lever, whether that is achieved using the left paw or the right paw. The demonstration of bilateral plasticity is important for this claim and, indeed, appears to be a characteristic of goal-directed action that differentiates from other forms of action-related learning, such as is induced by sensory-motor habits or skills (Kent et al., 2013; Xiong et al., 2015).

### The BLA-pDMS pathway does not undergo plasticity during goal-directed learning

We found, with pathway specific toxicogenetic ablation, that the BLA-pDMS was not required for the acquisition of goal-directed action, and that the pathway did not undergo long-term plasticity as a result of training. It remains possible that functional plasticity in the BLA-pDMS pathway exists on to striatal interneurons or only mediates short-term changes in activity, perhaps for instrumental performance rather than learning. Interestingly, a prior study (Corbit et al., 2013) found evidence that BLA-pDMS disconnection affects choice performance between previously acquired goal-directed actions, something that the PL and so, presumably, the BLA-PL pathway, does not. However, in the Corbit et al. (2013) study, rats received unilateral lesions of the BLA and contralateral (disconnection) lesions (or ipsilateral control) of the pDMS. As such, it is possible, as we found here with regard to the PL, that a third structure mediates this performance related interaction of BLA and pDMS, perhaps the medial orbitofrontal cortex (Bradfield et al., 2015).

An interesting observation in the present results is that the BLA-pDMS pathway appeared to be biased towards iSPNs. Nearly every iSPN recorded (93%) was found to have active input from the BLA, in comparison to approximately 60% of dSPNs. Thus, it is possible that an important aspect of the BLA-pDMS projection is the modulation of iSPNs, something that recent findings from our lab suggest could be of more direct importance to updating goal-directed learning (Matamales et al., 2020).

### BLA-PL pathway is critical for both goal-directed learning and PL-pDMS plasticity

Perhaps the most intriguing finding in this series of experiments is the importance of the BLA-PL pathway to goal-directed learning and PL-pDMS plasticity. The bilateral ablation of this pathway produced insensitivity to outcome devaluation after goal-directed learning, something that was not observed in a previous report in which the BLA and PL were disconnected using asymmetrical lesions (Coutureau et al., 2009). Additionally, when BLA neurons were inhibited during the task using DREADDs, potentiation of the PL-pDMS pathway was abolished. These findings suggest that the BLA plays a direct modulatory role in the development of PL-pDMS plasticity during learning.

This finding is important because, based on the finding that projections from the BLA to the DMS spatially overlap with those from the PL (Mcdonald, 1991) - and, indeed, it has been reported that the vast majority (90%) of DMS cells received both PL and BLA inputs (Ma et al., 2017) – it might be supposed that the BLA involvement is direct to pDMS rather than indirect via PL. Furthermore, Ma et al. (2017) also reported that paired activation of PL and BLA afferents induced a supralinear integration, and that brief high frequency pairings (2 s at 50 Hz) induced potentiation of the PL-evoked local field potential whereas potentiation was not found with stimulation of BLA afferents alone. As such, there are good reasons to predict that the direct BLA input to the pDMS will be necessary for goal-directed learning. It was surprising, therefore, not only to find no evidence of plasticity in this projection on either dSPN or iSPNs, but that ablation of this pathway had no effect on goal-directed learning. The latter finding also counters an alternative explanation both for the failure to see potentiation of BLA input and for the effect of BLA inhibition on PL-pDMS plasticity; i.e., that the BLA serves to potentiate the PL input through an additional excitatory input. Were this the case then BLA-pDMS pathway ablation should have abolished goal-directed learning and it did not.

### Summary

In summary, this study demonstrates that the bilaterally-projecting PL projection to the pDMS is potentiated during goal-directed learning, specifically the projection to dSPNs. Furthermore, our findings indicate that the BLA-PL pathway is critical for this learning, and that the BLA provides information to the PL during the acquisition of goal-directed actions necessary for learning-related plasticity in the PL-pDMS pathway, and necessary for goal-directed learning. Together these results reconcile prior findings and elucidate the roles of key brain regions required for encoding the specific action-outcome associations necessary to acquire new goal-directed actions.

## Methods

### Lead contact and material availability

This study did not generate new unique reagents. Further information and requests for resources and reagents should be directed to and will be fulfilled by the Lead Contact, Bernard Balleine.

### Subjects

Male and female, 7–10 weeks old, C57Bl/6J–Quackenbush hybrid transgenic mice carrying bacterial artificial chromosome (BAC) that expresses enhanced green fluorescent protein (BAC-EGFP) under the control of the D2R promoter (Drd2-EGFP) were used for the majority of experiments. Mice were hemizygous. In experiments described in Fig. 1 and Fig. 5, C57BL/6 mice were used. Littermates were randomly assigned to experimental groups, and were all group housed. The mice were housed in a 12 h light/12 h dark cycle in a temperature-controlled (21°C) and humidity-controlled (50%) environment. Mice were food restricted to 85% of their projected free feeding weight throughout behavioral sessions, starting three days prior. All groups were composed of mixed male and female subjects. Statistical tests for sex differences were conducted throughout the study but at no point did male and female subjects differ whether in behavioral, immunofluorescence or electrophysiological measures.

All experiments were conducted according to the ethical guidelines approved by the University of New South Wales Animal Care & Ethics Committee.

### Surgeries

All surgeries were performed under aseptic conditions while mice were anaesthetized with isoflurane (5% induction, 0.5%–1.5% during) in a stereotactic frame (Kopf 940). Local anesthetic (Bupivacaine, 0.1 ml) was administered subcutaneously, and Carprieve (10 mg/kg, subcutaneous) was used for postoperative analgesia. All viral injections were performed using a Nanoject III (Drummond Scientific Company) at 1 nL/min. The injection pipette was held in place for 5 minutes after completion of virus delivery, and a further 2 minutes after being withdrawn 0.05 mm. Mice recovered for 18 – 25 days before experiments started. All viruses were obtained from UNC Vector Core or Addgene.

To assess the PL-pDMS pathway (Fig. 1, Fig. 4), we injected 220 nl of adeno-associated virus serotype 5 (AAV5) carrying channelrhodopsin (ChR2) under the control of the CaMKIIa promoter and tagged with the mCherry fluorophore (rAAV5/CaMKIIa-hChR2(H134R)-mCherry). The virus was injected unilaterally into the prelimbic area at coordinates: +1.65 mm AP, 0.4 mm ML, measured from bregma on the skull surface, and 1.4 mm DV from the brain surface. Virus infection spread in the PL is illustrated in Supp. Fig. 1.

To assess the BLA-pDMS pathway (Fig. 2), we injected 250 nl of the AAV5 virus described above, unilaterally into the BLA at coordinates: −1.2 mm AP, 3.3 mm ML, measured from bregma on the skull surface, and 4.4 mm DV from the brain surface.

For the Caspase lesion experiments outlined in Fig. 3, C57BL/6 mice were injected bilaterally with 220 nl of AAV.hSyn.HI.eGFP-Cre.WPRE.SV40 in either the PL or the pDMS, and 250 nl of AAV-flex-taCasp3-TEVp or PBS as vehicle control in the BLA. PL and BLA injections were as described above. pDMS coordinates were: −0.2 mm AP, 1.8 mm ML, measured from bregma on the skull surface, and 1.8 mm DV from the brain surface.

### Behavior

Med Associates operant chambers were used for instrumental conditioning. Initially, two sessions of magazine entry training were given, during which 30 grain pellets (20 mg each) were delivered on a random time 60 s schedule. Next, mice completed four instrumental training sessions in which they had to press a lever to receive a grain pellet. Each session lasted until either 30 grain pellets were delivered or 60 minutes passed. In the first continuous reinforcement (CRF) session, pellets were delivered on every lever press. In the following three sessions pellets were delivered on a random interval (RI) schedule of 15, 30 and 30 seconds respectively. Yoked control mice experienced the same training sessions, although essentially received random pellet delivery, of the same quantity to trained animals in each session, but non-contingent on lever presses.

Outcome devaluation testing was performed in the experiment relating to Figure 1, and followed previously published protocol (Shan et al., 2014). Mice were given a specific satiety devaluation test the day after the final training session. Half of the mice (pellet condition) were given 1 hour free access to grain pellets (“devalued” condition), which they normally earned during the training sessions, whereas the remainder (chow condition) were given free access to the chow that they received in their home cages, for the same duration as a control for the effects of general satiety (“other” condition). The devaluation effect was tested in a 5 min extinction test conducted immediately after the satiety treatment. No pellets were delivered during the test. A second devaluation test was conducted the day after the first devaluation test for which the satiety (pellet or chow) conditions were reversed.

The number of magazine entries per minute was calculated across sessions to give a measure of task concentration and adherence. Additionally, the mean reward collection time was calculated for each session, which is defined as the time between reward delivery and the mouse nose poking for the grain pellet.

The plasticity results found were due to the difference in learning experiences between mice that had contingent and noncontingent outcomes on lever press. Completely naive controls were not used as they previously showed no difference to yoked animals in gross plasticity due to this training (Shan et al., 2014).

### Brain slice preparation

Approximately 24 hours following the final instrumental conditioning session, mice were processed for *ex vivo* slice recording. Mice were briefly anesthetized with isoflurane and then perfused with ice-cold NMDG solution containing (in mM): 92 NMDG, 2.5 KCl, 1.6 NaH_2_PO_4_.2H_2_0, 30 NaHCO_3_, 20 HEPES, 25 glucose, 2 thiourea, 5 Na-ascorbate, 3 Na-pyruvate, 0.3 CaCl_2_, 10 MgSO_4_,7H_2_0 (saturated with 95% O_2_ and 5% CO_2_, pH 7.3-7.4, osmolarity 300–310 mOsm). Coronal slices (300 um) including the pDMS (approximately 0.2 to −0.6 around bregma) were cut on a vibratome (Leica Microsystems VT1200S) in the same ice-cold NMDG solution. Slices were recovered for approximately 12 minutes in the NMDG solution at 33ºC, and then transferred to room temperature ACSF solution containing: 126 NaCl, 2.5 KCl, 2 CaCl_2_, 2 MgSO_4_.7H_2_0, 1.2 NaH_2_PO_4_.2H_2_0, 25 NaHCO_3_, 11.1 glucose (saturated with 95% O_2_ and 5% CO_2_, pH 7.3-7.4, osmolarity 300–310 mOsm). Slices were left for at least an hour before recording.

### Electrophysiology

Slices were transferred to a recording chamber and neurons visualized under an upright microscope (BX51WI, Olympus) using differential interference contrast (DIC) Dodt tube optics, and superfused continuously (1.5 ml/min) with carboxygenated physiological ACSF at 33ºC. In voltage-clamp experiments Picrotoxin (100 uM) was added to the ACSF solution to suppress GABAergic currents. Whole-cell voltage-clamp recordings were made using electrodes (2–5 MΩ) containing internal solution (in mM): 120 cesium methanesulfonate, 15 CsCl, 8 NaCl, 10 HEPES, 0.4 EGTA, 3 QX-314, 2 Mg-ATP, and 0.3 Na-GTP (pH 7.3, osmolarity 285-290 mOsm/L). Current-clamp recordings (Fig. 4F) were made with an internal solution containing (in mM): 145 K-Gluconate, 10 HEPES, 1 EGTA, 2 MgCl_2_, 2 Mg-ATP, 0.3 Na_2_-GTP (pH 7.3, osmolarity 285-290 mOsm/L). Biocytin (0.1%) was added to internal solutions for labeling the sampled neurons during recording. Data acquisition was performed with an Axon Axopatch 200B amplifier (Molecular Devices), connected to a computer via an Instrutech ITC-18 interface. Liquid junction potentials were estimated at −10 mV and were not corrected. Somatic access resistance <25 MOhms was monitored and cells with unstable access resistance (>30% change) were discarded.

To activate axons containing ChR2 we generated 470 nm light with a collimated LED (Thorlabs M470L3-C1), which was intensity modulated with a T-Cube LED driver (ThorLabs) and passed through a 40× objective lens. Light pulse (5 ms) intensity was set to elicit a reliable postsynaptic current between 100 and 1000 pA. To record AMPA/NMDA plasticity measures, neurons were voltage-clamped either at −70 mV or at 40 mV. After a 5 minute stabilization period, EPSCs were evoked for 3-5 minutes at each membrane potential, with an inter-pulse interval of 20 seconds. Following these measures, paired-pulse stimulations were performed at −70 mV with a 50 ms interval.

The AMPA/NMDA ratio was calculated by dividing the mean peak amplitude of the EPSCs recorded at −70 mV (AMPA) by the averaged magnitude, 50–60ms after the stimulation artifact, of the EPSCs recorded at 40 mV (NMDA). Paired-pulse responses were averaged and the ratio of the second pulse to the first was calculated as the paired-pulse ratio. All analysis was performed with custom scripts in MATLAB.

### Cell identification

Cells for *ex vivo* recording were identified primarily by their GFP expression: GFP positive cells in the DMS of Drd2-EGFP mice were recorded as dopamine D2 receptor positive, indirect pathway SPNs. GFP negative cells, of a similar shape, were recorded as putative D1R-expressing, direct pathway SPNs. The GFP expression profile of Drd2-EGFP mice has been extensively validated in previous studies (Bertran-Gonzalez et al., 2008; Matamales et al., 2009).

Immediately after recording, brain slices containing biocytin-filled neurons were fixed overnight in 4% paraformaldehyde/0.1 M phosphate buffer (PB) solution. Slices were rinsed and then incubated in Alexa Fluor 647-conjugated Streptavidin (1:1000; Life Technologies) for 2 h at room temperature. Stained slices were rinsed, mounted onto glass slides, dried, and coverslipped with Vectashield mounting medium (Vector Laboratories). Images were obtained using laser scanning confocal microscopy (Fluoview FV1000, BX61WI microscope, Olympus) and a spinning disk field scanning confocal system (Nikon Ti-E with Andor Diskovery system). Recorded cells that were successfully filled with biocytin and recovered were verified to have spines on their dendrites, indicative of a SPN and ruling out the vast majority of interneurons, which are predominantly aspiny (Tepper et al., 2010).

### Histology

Following brain slicing for electrophysiology, brain chunks containing viral injection sites were fixed overnight in 4% paraformaldehyde/0.1 M phosphate buffer solution. Slices (40 um) were made, then rinsed, mounted onto glass slides, dried, and coverslipped with Vectashield mounting medium (Vector Laboratories). A sampling of images throughout the viral infection spread were obtained using the spinning disk field scanning confocal system previously described.

### Quantification and statistical analysis

All data are shown as mean ± SEM unless otherwise specified. Sample sizes (n) are defined as the number of cells, and the number of animals used is also stated in the Figure legends. Comparisons were conducted with two or three-way ANOVAs, or t tests, as described in Figure legends, using R 3.6 (the ‘car’ library) or Prism 7 (GraphPad). ANOVAs were followed by Tukey’s post-tests for multiple comparisons where appropriate. P < 0.05 was considered statistically significant.

### Data and code availability

MATLAB code used to analyze AMPA/NMDA ratio and paired-pulse recordings is available at the following GitHub repository: https://github.com/simonfisher/goal-directed-plasticity

## Acknowledgments

The research reported in this manuscript was supported by a Discovery Grant from the Australian Research Council, #DP150104878, and a both Project Grants, GNT #1089252, #1165346, and a Senior Principal Research Fellowship, #1079561, from the National Health and Medical Research Council of Australia to BWB.

## Author Contributions

S.F. conducted behavioral and electrophysiology experiments; L.F. conducted behavioral and Caspase lesion experiments, and performed image analysis; S.F., B.B., and J.B-G designed the experiments and wrote the paper.

## Supplemental Information

**Figure S1.**
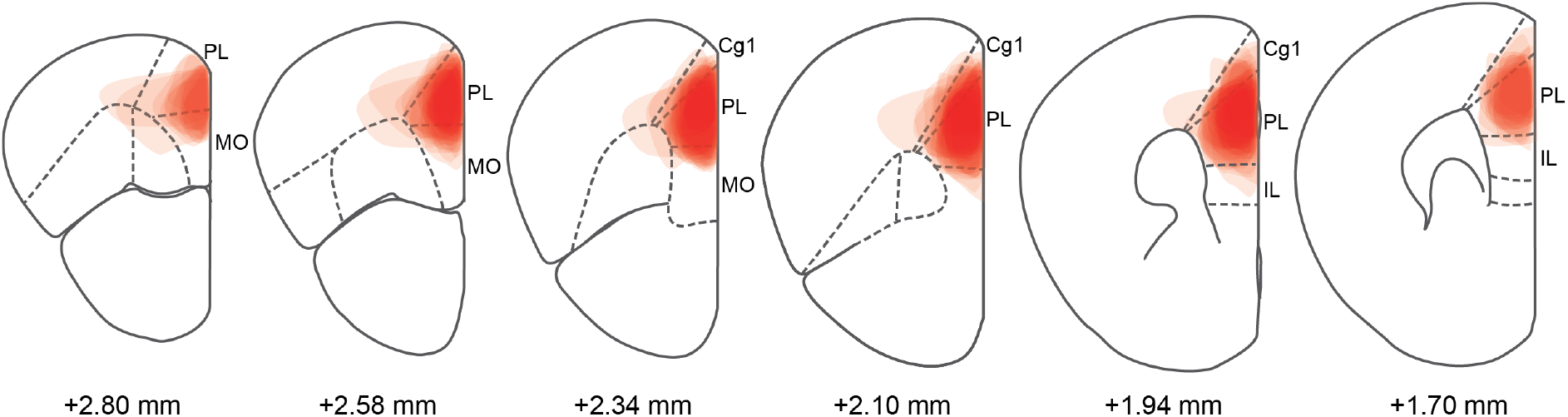
Prelimbic injection sites relating to experiments described in Figure 1. Each translucent region corresponds to the spread of the viral infection for individual mice. Coordinates below coronal slice diagrams relate to position anterior to Bregma. PL = prelimbic cortex; MO = medial orbital cortex; Cg1 = cingulate cortex area 1; IL = infralimbic cortex.

**Figure S2.**
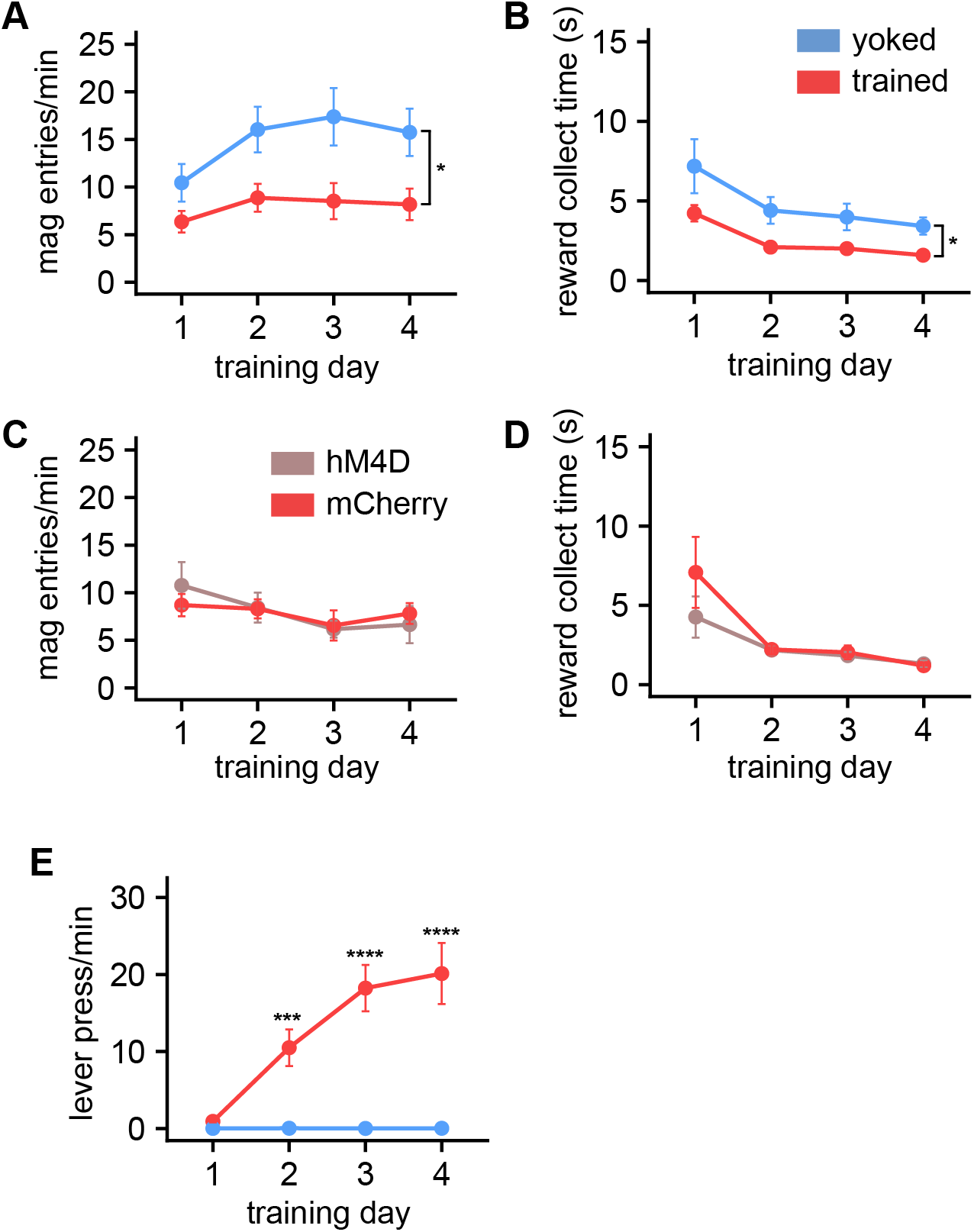
Behavioral data relating Figures 1 and 4. **(A)** yoked control mice had a significantly higher rate of magazine entries across the training sessions (two-way RM ANOVA, F1,30 = 6.35, P = 0.02). (**B)** trained mice exhibited a significantly lower time to collect the food pellet reward, as measured from the time of reward delivery, than yoked control mice (two-way RM ANOVA, F1,30 = 6.84, P = 0.01). (**C)** no difference was found between trained animals with active hM4D DREADD and trained mCherry controls in magazine entries per minute or (**D)** reward collection time. **(E)** Learning behavioral results of animals in the PL contralateral group. Mice in the trained group, with reward delivery contingent on lever presses, had a significantly higher press rate than yoked controls for which reward delivery was non-contingent (two-way RM ANOVA: session x group, F_(3,57)_ = 32.53, *P* < 0.0001; post hoc tests, \*\*\**P* = 0.0001, \*\*\*\**P* < 0.0001).

**Figure S3.**
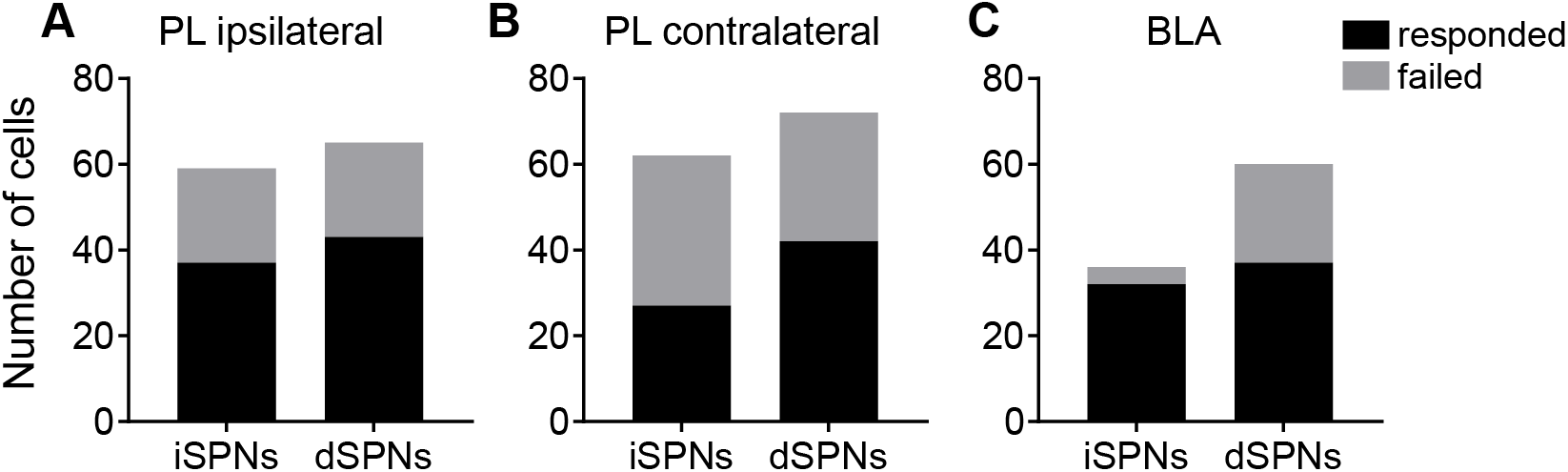
Proportion of dSPNs responded to optical stimulation in the ipsilateral and contralateral pathways. **(A)** The proportion of cells, during stable and healthy whole cell recordings, that responded to optical stimulation of PL ipsilateral, **(B)** PL contralateral, and **(C)** BLA ipsilateral projections.

**Figure S4.**
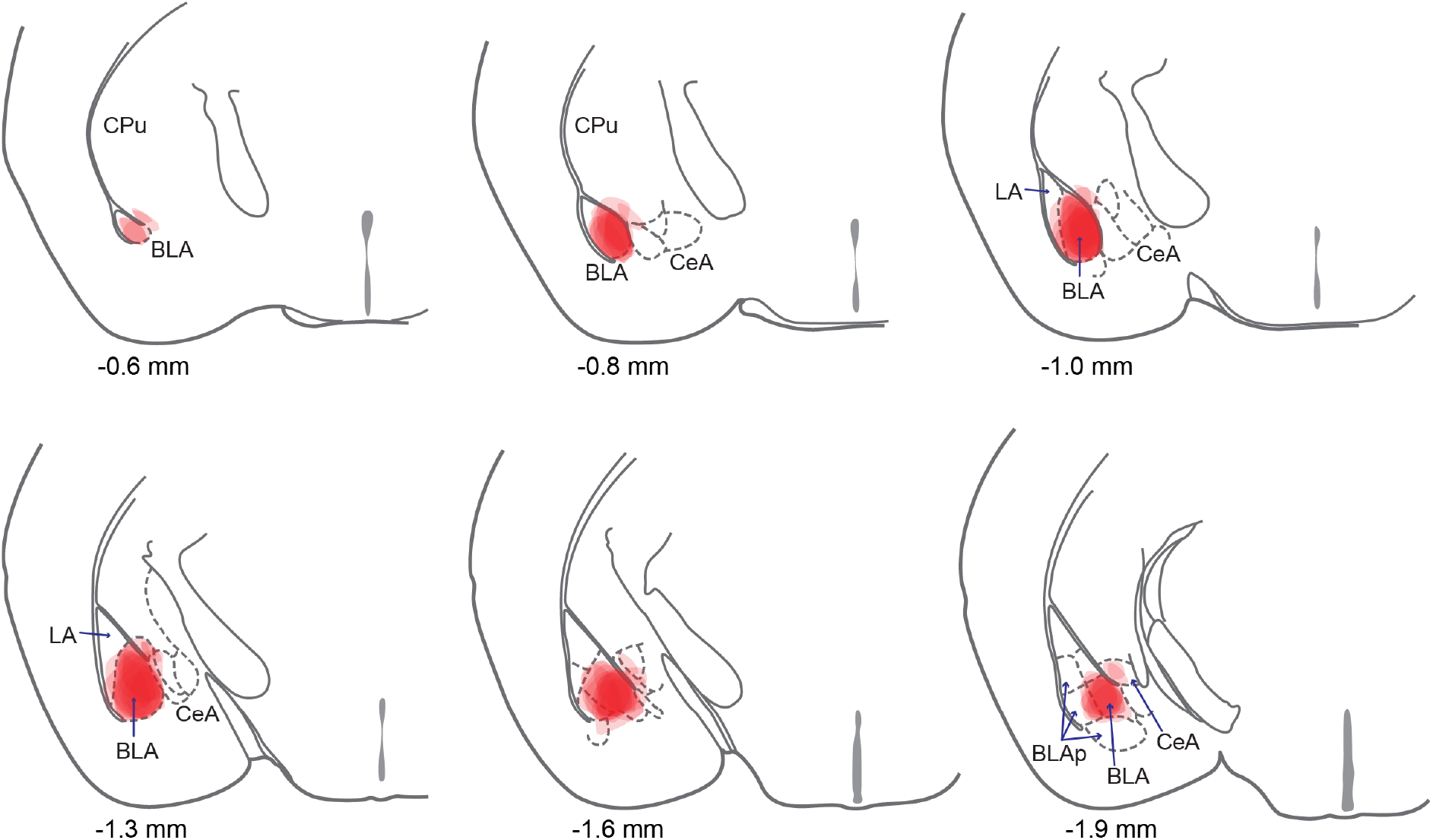
Basolateral amygdala injection sites relating to the study described in Figure 2. Each translucent region corresponds to the spread of cell infection. Coordinates below coronal slice diagrams relate to the position posterior to Bregma. CPu = caudate putamen/striatum; BLA = basolateral amygdala; CeA = central amygdala nuclei; LA = lateral amygdala; BLAp = posterior portion of the BLA.

**Figure S5.**
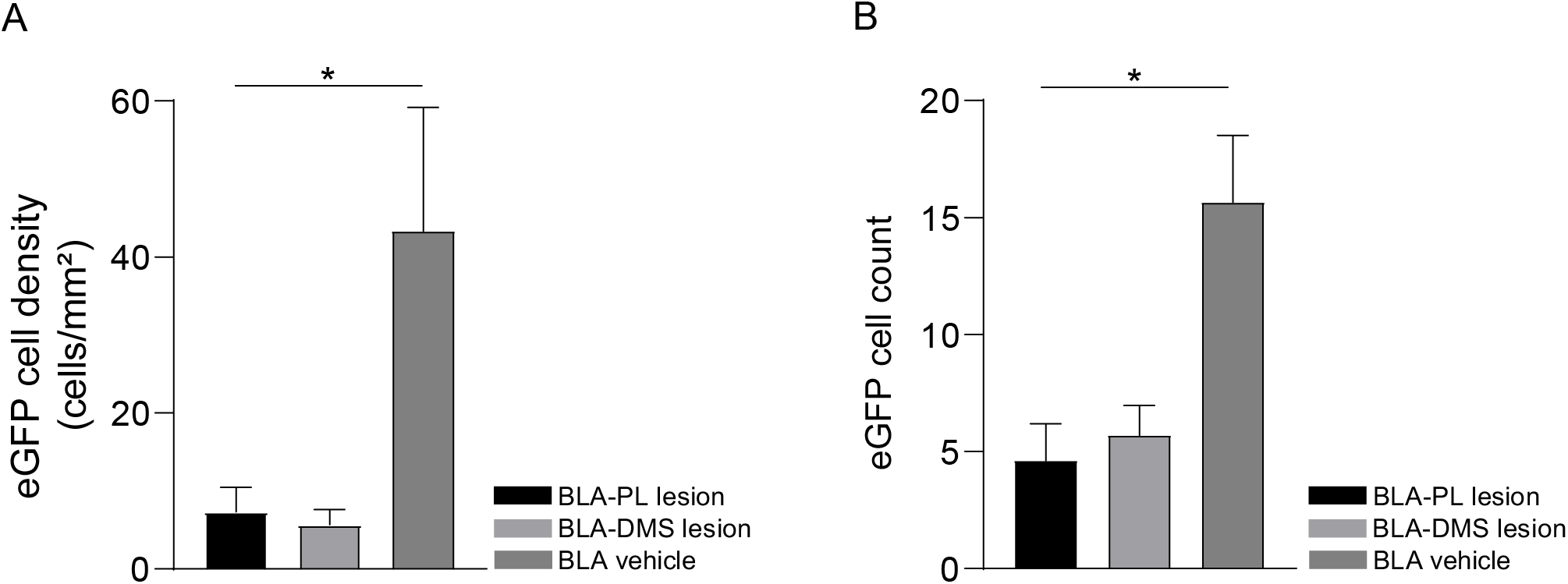
BLA Caspase lesion validation, relating to Fig. 3. **(A)** Quantification of eGFP cell density in the BLA between groups. There was significant loss of eGFP+ve cell density in the BLA-PL lesion group (p=0.0322) and BLA-DMS lesion group (p=0.0247) relative to the BLA vehicle control (F (2, 21) = 5.206, P=0.0146, one-way ANOVA with Tukey’s post-hoc comparisons). **(B)** Quantification of eGFP cell count in the BLA between groups. There was significant loss of eGFP+ve cells in the BLA-PL lesion group (0.0005) and BLA-DMS lesion group (p=0.0020) relative to the BLA vehicle control (F (2, 141) = 8.996, P=0.0002), one-way ANOVA).

**Figure S6.**
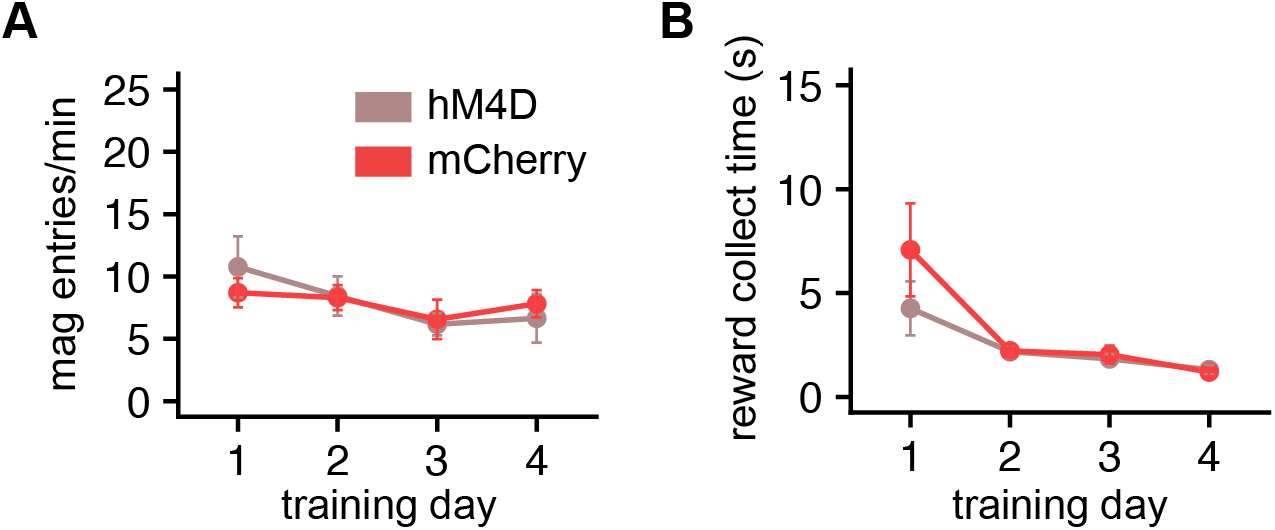
Behavioral effect of DREADD-induced inactivation of BLA. **(A)** There was no difference between hM4D-expressing mice and mCherry control mice in magazine entries across the training sessions, nor in **(B)** time to collect the food pellet reward, as measured from the time of reward delivery.

